# Protein structure search to support the development of protein structure prediction methods

**DOI:** 10.1101/2020.06.03.131821

**Authors:** Ronald Ayoub, Yugyung Lee

## Abstract

Protein structure prediction is a long-standing unsolved problem in molecular biology that has seen renewed interest with the recent success of deep learning with AlphaFold at CASP13. While developing and evaluating protein structure prediction methods, researchers may want to identify the most similar known structures to their predicted structures. These predicted structures often have low sequence and structure similarity to known structures. We show how RUPEE, a purely geometric protein structure search, is able to identify the structures most similar to structure predictions, regardless of how they vary from known structures, something existing protein structure searches struggle with. RUPEE accomplishes this through the use of a novel linear encoding of protein structures as a sequence of residue descriptors. Using a fast Needleman-Wunsch algorithm, RUPEE is able to perform alignments on the sequences of residue descriptors for every available structure. This is followed by a series of increasingly accurate structure alignments from TM-align alignments initialized with the Needleman-Wunsch residue descriptor alignments to standard TM-align alignments of the final results. By using alignment normalization effectively at each stage, RUPEE also can execute containment searches in addition to full-length searches to identify structural motifs within proteins. We compare the results of RUPEE to mTM-align, SSM, CATHEDRAL and VAST using a benchmark derived from the protein structure predictions submitted to CASP13. RUPEE identifies better alignments on average with respect to RMSD and TM-score as well as Q-score and SSAP-score, scores specific to SSM and CATHEDRAL, respectively. Finally, we show a sample of the top-scoring alignments that RUPEE identified that none of the other protein structure searches we compared to were able to identify.

The RUPEE protein structure search is available at https://ayoubresearch.com. Code and data are available at https://github.com/rayoub/rupee.

## Introduction

Determining the structure of a protein is an important step toward understanding its function. There are approximately 150,000 solved protein structures currently stored in the protien data bank (PDB) [1], the global repository for experimentally determined protein structures. On the other hand, UniProt [2], the universal protein knowledgebase, currently provides over 60 million protein sequences. From this, it is apparent that protein structure determination is moving at a slower pace than protein sequencing and may be serving as a bottleneck in a variety of research efforts from protein design to drug discovery. Being able to predict a protein structure from its amino acid sequence would address this problem. However, protein structure prediction remains a central unsolved problem in molecular biology [3].

CASP is a biannual blind competition for protein structure prediction that began in 1994 [4]. Progress had been slow until the success of coevolutionary methods in contact prediction demonstrated in CASP11 [5]. The recent success of AlphaFold at CASP13 [3] using deep learning combined with coevolutionary methods has renewed interest in the problem. While AlphaFold’s performance was remarkable, it depends on the availability of sufficiently large multiple sequence alignments (MSA) for its use of coevolutionary methods, which may not be available for all target structures. Even when a large enough MSA is available, AlphaFold as well as traditional physics-based approaches to protein structure prediction, such as Rosetta [6], have not reached the desired level of accuracy [3].

Previously, we introduced RUPEE [7], a purely geometric protein structure search with no dependence on sequences or clustering. We compared our results with the mTM-align structure search [8], the secondary structure matching (SSM) search [9], and the CATHEDRAL structural scan [10], and found RUPEE is equal to or better than some of the best available structure searches on a benchmark of known protein structures. Additionally, we showed RUPEE, on average, returns results faster than mTM-align and CATHEDRAL.

Since the release of RUPEE [7], we have observed that RUPEE has been used to upload protein structures that were the output of a protein structure prediction method in order to identify the most similar known structure to the predicted structures. For the most part, these uploaded protein structures have had low sequence and structure similarity to known structures in the PDB. This low similarity to known structures is to be expected given the limited accuracy of current protein structure prediction methods.

When searching for structures with low sequence and structure similarity to known structures, the importance of small differences in structure similarity becomes proportionally larger since they comprise a larger percentage of the overall similarity. Protein structure searches that rely on sequences or clustering may miss these small differences in similarity because they do not consider every structure individually. Moreover, with respect to the use of sequences, while high sequence similarity usually indicates high structure similarity [11], high structure similarity has been observed even for structures with low sequence similarity since structure is more conserved in evolution than sequence [12]. While sequences or clustering can be used to reduce the number of structures that have to be examined, thereby decreasing response times, RUPEE can examine every structure individually during a search, ensuring all similar structures are found, and does so in a reasonable amount of time.

While the original top-aligned search mode for RUPEE [7] has no dependence on sequences or clustering, it often lacks sufficient sensitivity to find the most similar match for a structure with low structure similarity to known structures. This lack of sensitivity for RUPEE top-aligned search mode with respect to low similarity searches is due to the lower accuracy of its initial structure similarity estimates used to filter candidate matches. Recognizing the need for a structure search with more sensitivity than top-aligned search mode and still having no dependence on sequences or clustering, we have added an additional search mode to RUPEE with increased sensitivity called all-aligned search mode.

Like our previous work on RUPEE [7], again we compare the results of RUPEE against mTM-align [8], SSM [9] and CATHEDRAL [10], but this time we do so for all-aligned search mode. Additionally, this time we also compare to the VAST protein structure search [13]. For comparisons, we use a new benchmark derived from protein structure predictions of free-modelling targets in CASP13 available at the CASP web site [14]. Previously we showed that RUPEE, in top-aligned search mode, is equal to or better than those we compared to for a benchmark of known protein structures. Here, we show that RUPEE, in all-aligned search mode, is better than those we compare to using a benchmark drawn from the output of protein structure prediction methods.

While it is possible to perform a protein structure search by exhaustively comparing to every available structure using pairwise structure alignments, we do not compare RUPEE to pairwise structure alignment tools used in this manner. If an exhaustive search is practical, it will always be optimal with respect to the pairwise structure alignment tool used. However, an exhaustive search may not be practical given the resources needed to achieve a desired response time. To illustrate this point, consider the number of pairwise structure alignments needed to perform a single search because the pairwise structure alignments dominate the response time for RUPEE. For searching whole PDB chains in the RUPEE database, presently containing roughly 440,000 structures, RUPEE performs only 8000 pairwise structure alignments. Given the same amount of resources, an exhaustive search will take roughly 440, 000/8000 = 55 times longer. With the number of structures available in the PDB growing rapidly, this difference will continue to increase. Therefore, we only compare to other protein structure searches which are designed, like RUPEE, to return quality results with much faster response times.

The protein structure searches we compare to represent a good mix of approaches. For structure searches that depend on sequences and clustering, mTM-align [8] is among the best available and is capable of handling uploaded structures with the same response-times as for searching on a structure id. SSM [9] is a good example of a fast graph-theoretic structure search with no dependence on sequences and clustering. However, SSM’s speed is at the expense of sensitivity since it depends on the spatial orientation and connectivity of secondary structure elements, which fails to capture the complexity of loops. CATHEDRAL [10] also uses a fast graph-theoretic approach that is more accurate than SSM, but still lacks sufficient sensitivity to identify the most similar structure matches for low similarity searches. Moreover, although the CATHEDRAL structural scan does not depend on clustering directly, it only searches on and returns results for representatives of sequence clusters at 35% similarity [15]. The VAST protein structure search [13] is similar to SSM in that it depends on the spatial orientation and connectivity of secondary structure elements. Although VAST is much slower than SSM for uploaded structures, its search database is more recent than that of SSM.

Previously, the RUPEE protein structure search focused on full-length matches. If you wanted to search for structures similar to a domain, you could search one of the protein domain classification databases such as SCOPe [16], CATH [15], or ECOD [17]. On the other hand, if you wanted to search for structures similar to a whole chain, you could search whole chains from the PDB. While the ability of RUPEE to search multiple databases for any given structure is flexible, we have extended this flexibility by introducing explicit search types for *Contained-In* and *Contains* searches in addition to the *Full-Length* search type. The Contained-In search type searches for structure that are contained in the query structure and the Contains search type searches for structures that the query structure contains. With support for containment searches, RUPEE can now be used to search for structural motifs within proteins.

Together, all-aligned search mode and containment searches are significant additions to RUPEE that support the development of protein structure prediction methods by allowing researchers to find the structures most similar to the output of their predictions, regardless of how they vary from known protein structures.

## Methods

We first give a brief outline of our linear encoding of protein structures described in more detail in our previous work on RUPEE [7], which still remains at the core of the RUPEE protein structure search. Then, we describe our approach to top-aligned to provide context followed by the addition of all-aligned.

### Linear encoding of protein structure

Previously [7], we introduced a linear encoding of protein structures based on torsion angle regions. We determined these regions by plotting a random sampling of torsion angles. Traditionally, the Ramachandran plot would be used for the plotting. However, we found that the discontinuities within the Ramachandran plot due to the cyclical nature of torsion angles made the Ramachandran plot poorly suited for this task. Instead, we plotted the sampled torsion angles on a polar plot we also introduced in [7] to clearly define continuous regions. Figure 1 shows our polar plot alongside its corresponding Ramachandran plot and Figure 2 shows the defined regions for helices, strands, and coil using the polar plot. We assigned an integer, referred to as a residue descriptor, to each defined region. The DSSP [18] secondary structure assignment codes for turns (‘T’) and bridges (‘B’) are assigned descriptors 11 and 12 respectively, regardless of region.

**Figure 1.**
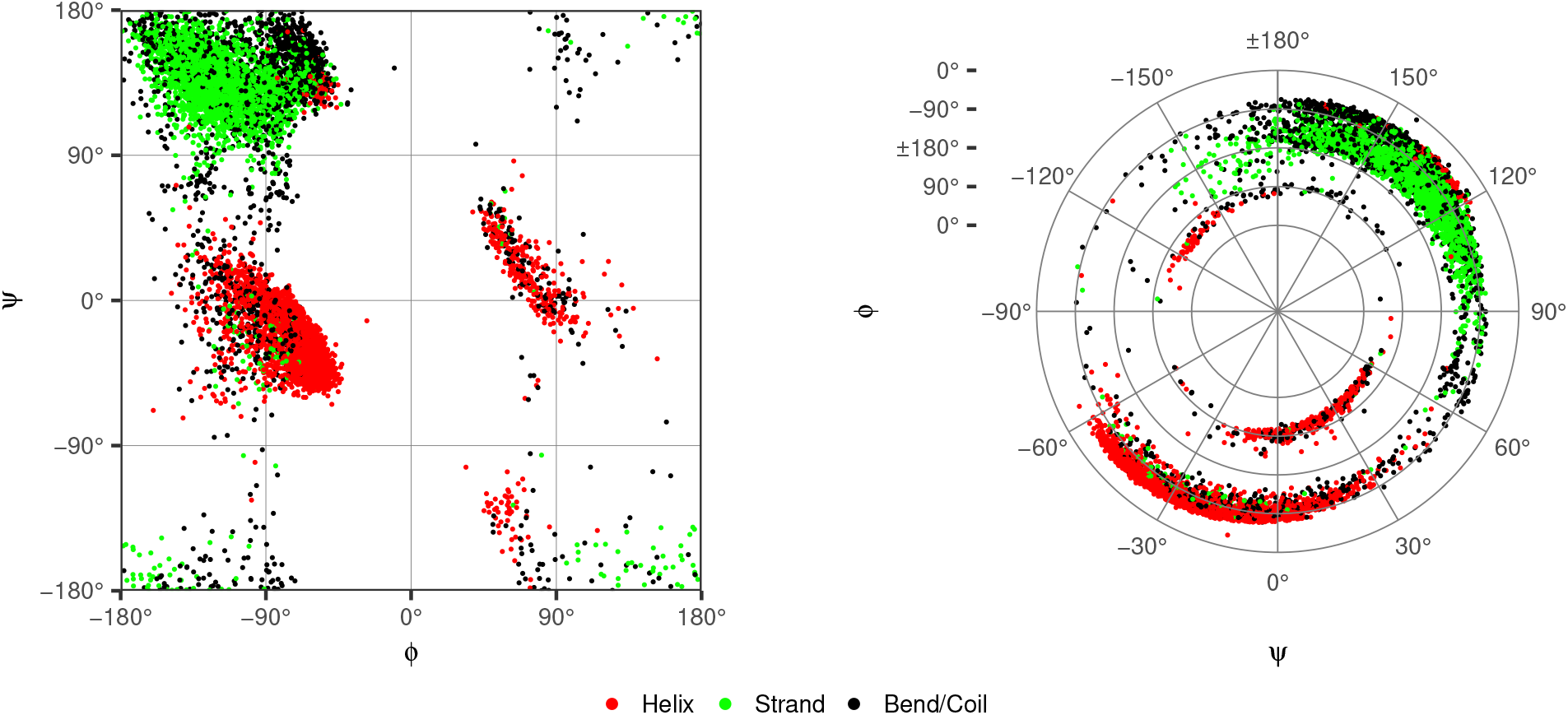
Ramachandran plot (right) and polar plot (left) of randomly sampled torsion angles

**Figure 2.**
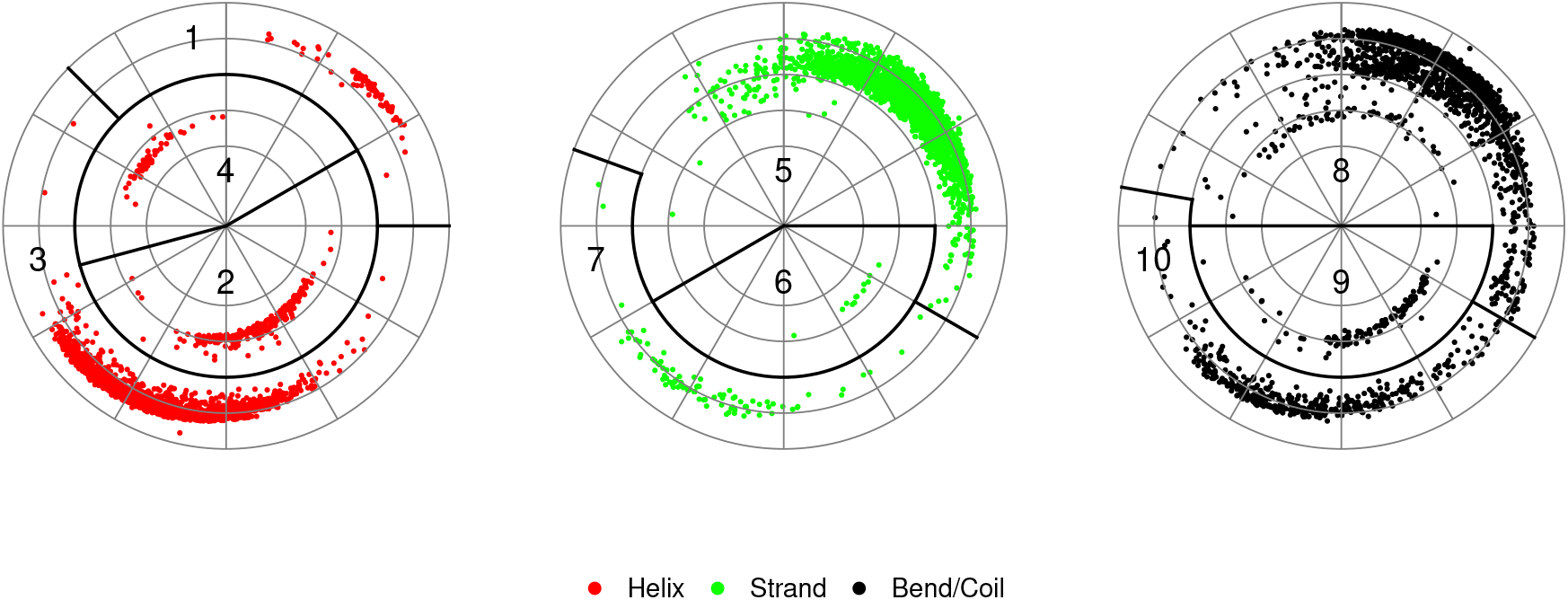
Polar plots of randomly sampled torsion angles with designated descriptors for region and secondary structure combinations

As an example of our linear encoding, Figure 3 shows a typical *β*-turn-*β* motif annotated with the residue descriptors corresponding to the sequence shown below. The underlined elements in (1) correspond to the underlined elements in (2), (4), and (5) below to help illustrate the subsequent transformations from descriptors to shingles and finally to hashes.

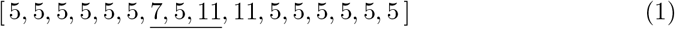

**Figure 3.**
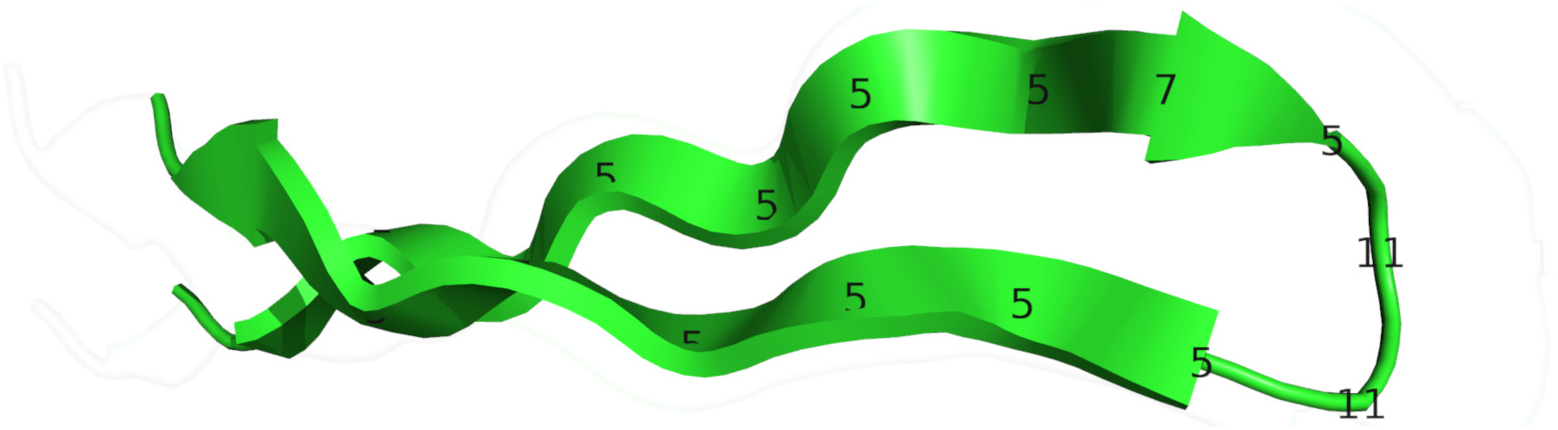
*β*-turn-*β* motif from CATH domain 1nycA00

Next, we derive a multiset of overlapping 3-grams of residue descriptors, where a 3-gram is three consecutive residue descriptors. This representation is often referred to as shingling, given their likeness to overlapping roofing shingles [19]. The overlap between shingles ensures some of the order information within the original sequence is preserved in the multiset.

By shingling, we obtain a multiset of ordered sequence from an ordered sequence of residue descriptors. As an example, the sequence in (1) becomes the following multiset of shingles.

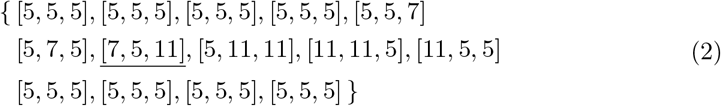

Next, each shingle *s* is hashed to an integer *s*_*hash*_ as shown in (3), where *s*_*i*_ is the *i*^th^ descriptor of the shingle *s*.

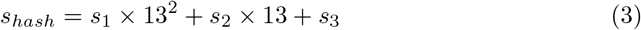

After hashing, the multiset in (2) becomes the following multiset of integers.

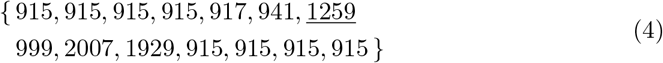

The hashing completes the transformation of an ordered sequence of residue descriptors to a multiset of integers that still retains some of the order information present in the original sequence.

In (4) the value 915, corresponding to the shingle [5, 5, 5], occurs frequently indicating the presence of *β*-strands. To address this lack of specificity, we introduced a heuristic we call *run position encoding* (RPE) [7], where a run is a consecutive sequence of identical descriptors. To distinguish between short and long runs, thereby increasing the specificity of the shingles, we add a factor of 10^5^ to each shingle hash as a function of the first residue’s position in a run. Applying RPE to the multiset of integers in (4) gives

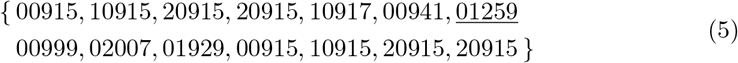

where the leading zero run factors are shown for clarity.

The pyramidal approach for the run factors used in RPE preserves matches at the boundaries between secondary structure runs and loops that would not otherwise be preserved in the presence of differences in run lengths of one or more.

### Search modes

Before RUPEE can service a search request, an offline process has to be executed in order to index the available protein structures. This index consist of residue coordinates, 3-grams, min-hashes and band-hashes for LSH banding. If a user searches on a structure id, its representation will already be stored in the index. On the other hand, if a user uploads a protein structure, it will be parsed into residue coordinates, 3-grams, min-hashes and band-hashes. Aside from the initial parsing, searches on uploaded structure are identical to searches by structure id. Figure 4 shows a flowchart for the RUPEE top-aligned and all-aligned search modes described below.

**Figure 4.**
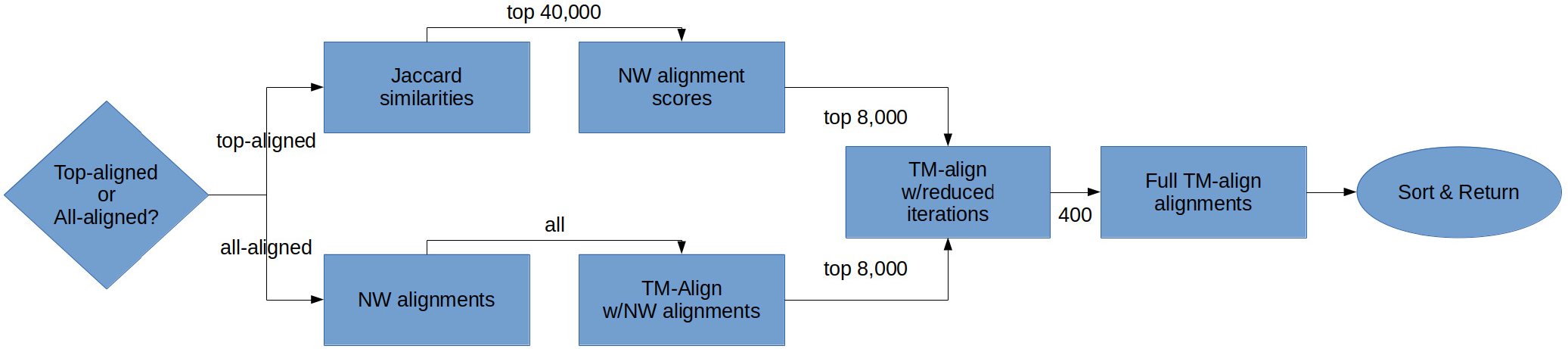
Flowchart for the RUPEE top-aligned and all-aligned search modes. It can be assumed that linear encodings and min-hashes for all structures other than the query structure have been stored via an offline indexing process and are accessible throughout the flowchart.

### Top-aligned search mode

When representing each protein structure using a multiset of integers as shown in (5), we define full-length similarity for a candidate pair of structures *a* and *b* using the Jaccard similarity for multisets [20],

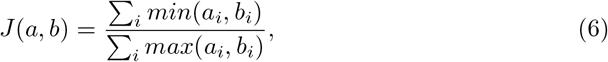

where *i* ranges over all possible shingle hashes *s*_*i*_ and *a*_*i*_ and *b*_*i*_ give the counts of shingle hash *s*_*i*_ in structures *a* and *b*, respectively.

For top-aligned, RUPEE uses min-hashing [21] and locality sensitive hashing (LSH) [22] as described in detail in our previous work [7] to quickly estimate the Jaccard similarity of a query protein against all available structures in the searched database. Then, the top-scoring 40000 protein structures are obtained based on the Jaccard similarity estimates to complete the initial filtering.

Next, regarding the multiset in (5) as an ordered sequence of integers, we obtain more accurate similarity scores for the filtered matches by performing global alignments using the residue descriptor sequences with a simplified Needleman-Wunsch (NW) [23] dynamic programming algorithm where mismatches and gaps are penalized −1 points and matches are awarded +1 points. For containment searches, depending on whether or not the search type is Contained-In or Contains, one of the sides of the dynamic programming matrix is not penalized for the opening gap and end gap. This is often referred to as semi-global sequence alignment [24].

After the NW algorithm is performed on the top 40000 protein structures from the initial filtering, the top-scoring 8000 structures are obtained for subsequent pairwise structure alignments. Pairwise structure alignment, the most accurate method for comparing protein structures, involves finding a set of spatial rotations and translations for two protein structures that minimizes a distance metric. Traditionally, the root mean squared deviation (RMSD) between *α*-carbons of aligned residues is minimized. However, the RMSD score does not factor in the distance between unaligned residues nor does it consider the percentage of aligned residues, that is, alignment *coverage*. RMSD scores also have some dependence on the length of the aligned proteins. On the other hand, the TM-score [25] takes all residues into account and normalizes for both coverage and length of the aligned proteins. For this reason, TM-score is frequently used in scoring protein structure alignments and accordingly, we use it for our alignment scoring.

RUPEE uses TM-align [26] for performing pairwise structure alignments. TM-align uses a rotation matrix designed to maximize the TM-score rather than minimizing the RMSD along with dynamic programming to find the best alignment. Similar to how we used global and semi-global sequence alignments with the NW algorithm for full-length and containment search types respectively, we apply the same logic to how normalization is used in the TM-align algorithm to be compatible with the NW algorithm. For Full-Length searches we normalize by the average length of both structures, for Contained-In searches we normalize by the query structure, and for Contains searches we normalize by the matched structures. The three types of normalization available in TM-align and how they affect searches are discussed in greater detail in our previous work on RUPEE [7].

When doing the pairwise alignments, we take the top-scoring 8000 structures from the NW alignment filter and perform structure alignments using TM-align with a reduced number of dynamic programming iterations. Next, we sort these structure alignments by TM-score and obtain the top-scoring 400 structures. Finally, we perform structure alignments using TM-align on the top-scoring 400 structures using the default number of dynamic programming iterations and return the results sorted by TM-score.

The filter sizes of 40000, 8000 and 400 have been chosen based on quality of results and speed. We have found that increasing the size of either of these filters results in only marginal improvements in the quality of results. Given that performing the structure alignments is the most time-consuming aspect of the RUPEE structure search, the marginal improvements gained from larger filter sizes have to be balanced against the number of structure alignments performed.

### All-aligned search mode

While top-aligned may be sufficient for searching for known protein structures [7], the need for greater sensitivity arises when searching with structure predictions that may only have a maximum TM-score of less than 0.50 when compared against all available structures. Furthermore, for top-aligned, the effectiveness of the containment searches is limited by the initial filtering using Jaccard similarity estimates, which biases the initial filtering toward full-length matches. We address both of these concerns with the addition of all-aligned.

In contrast to top-aligned, all-aligned skips the initial step of using min-hashing and LSH filtering. Instead, all-aligned runs the NW algorithm on all available structures to obtain the residue descriptor sequence alignments rather than the NW alignment scores as in top-aligned. The residue descriptor sequence alignments are then passed into TM-align as the initial alignments and TM-align is set to stick to those initial alignments. Skipping the min-hashing and LSH filtering combined with initializing TM-align with the NW sequence alignments increases the sensitivity of RUPEE in all-aligned and can support containment searches at all stages of processing.

Once the TM-align algorithm is run on all available structures using the NW residue descriptor alignments as initial alignments, the top-scoring 8000 are obtained. As done in top-aligned, we run TM-align with a reduced number of iterations on these 8000 to obtain the top-scoring 400 and finally run TM-align with the default number of iterations on these to obtain the final results.

While running NW sequence alignments on all available structures reduces the scalability of all-aligned, it is still reasonably fast. With the current count of structures in the PDB still being less than a million, estimated alignments can be calculated on the entire collection in less than 10 minutes on structures with 400 or fewer residues.

## Results

Like our previous work on RUPEE [7], we compare the average scores of ranked results to those of mTM-align [8], SSM [9], CATHEDRAL [10] and VAST [13]. However, this time we compare results for both RUPEE all-aligned and top-aligned search modes and instead of using a benchmark of known protein structures we use a benchmark derived from structure predictions submitted to CASP13. We perform pairwise comparisons with each structure search individually to reduce sources of systemic error in our evaluation. Each of these pairwise comparisons to mTM-align, SSM, CATHEDRAL and VAST is discussed in its own section below. We also compare RUPEE all-aligned to RUPEE top-aligned using the benchmark of predicted protein structures and a benchmark of known protein structures. After comparing results, we provide a sample of the top-scoring alignments RUPEE was able to identify that mTM-align, SSM, CATHEDRAL and VAST all failed to identify.

To evaluate the results of RUPEE against mTM-align [8], SSM [9], CATHEDRAL [10] and VAST [13] for the case of providing support for the development of protein structure prediction methods, we derived our initial benchmark from structure predictions submitted to CASP13. To ensure the benchmark was challenging, we only considered predictions submitted for the 25 single-segment free-modeling (FM) target domains in CASP13 [14]. To ensure the benchmark was not too challenging, we only considered the first designated predictions of the top 10 performing groups ranked by the Assessors’ formula (GDT_TS + QCS) applied to free-modeling targets. We call this benchmark casp_d250 since it consists of 250 structures, corresponding to 25 target domains for each of the 10 top-performing groups. The top 10 performing CASP13 prediction groups are shown in Table 1.

**Table 1.** CASP13 Prediction Groups.

| Number | Name |
| --- | --- |
| 043 | A7D |
| 322 | Zhang |
| 145 | QUARK |
| 089 | MULTICOM |
| 261 | Zhang-Server |
| 224 | Destini |
| 498 | RaptorX-Contact |
| 197 | MESHI |
| 354 | wfAll-Cheng |
| 196 | Grudinin |

Top 10 performing CASP13 prediction groups ranked by the Assessors’ formula (GDT_TS + QCS) applied to free-modeling targets.

The difficulty of the casp_d250 benchmark presented challenges for some protein structure searches. While we would have liked to compare our results to DALI [27], another popular purely geometric protein structure search based on inter-residue distances, DALI was unable to return any results for a majority of the benchmark structures. On the other hand, mTM-align, SSM, CATHEDRAL and VAST returned results for all but a few missed structures. To avoid comparing against potential bugs, when there are missed structures for a specific comparison, we create a new benchmark for that comparison that excludes the missed structures from our initial casp_d250 benchmark. For all benchmarks structures, we required that at least 100 results are returned, with the exception of mTM-align, which only returned 10 results for a majority of structures. The benchmark names indicate the number of structures included in the benchmark and appear in the title of each plot below.

All benchmark definitions can be found in S1 Benchmarks. We uniquely identify each benchmark structure using the format 〈target〉TS〈group〉-〈domain〉. For instance, the prediction submitted by the AlphaFold team named A7D and numbered 043 for the second domain in the target T0960 is referred to as T0960TS043-D2.

### Scoring

In order to evaluate scoring fairly, we took a number of measures common to all comparisons below. First, in addition to the search types *Contained-In*, *Contains* and *Full-Length* that all depend on the TM-score as described above, we have added the additional search types *Q-score* [9] and *SSAP-score* [28] to RUPEE in order to perform comparisons to SSM and CATHEDRAL using their native scores, respectively. We also have added the *RMSD* search type to RUPEE for its general usefulness and ubiquity. The additional search types demonstrate the pluggable nature of RUPEE. Although we still use TM-align for all internal pairwise alignments after the initial filtering and NW alignments, we are able to easily apply different scores to the resultant alignments provided by TM-align besides the TM-score.

Given that we use a number of different scores for evaluations that may differ from the native scoring used by a structure search, we take their top 100 results based on their native scoring and re-sort them based on the compared score before making comparisons. If we did not re-sort by the compared score, it would allow poor scoring structures based on the compared score to remain at higher ranks and bias the results favorably for RUPEE. Due to this re-sorting, we are careful not to draw conclusions regarding scoring at a particular rank but instead consider the top 100 results as an aggregate to be evaluated.

For some comparisons below, we compare the RUPEE structure search to a protein structure search using a score that the latter may not have a corresponding search type for. For example, CATHEDRAL does not have an option to search or sort by RMSD. In our comparison to CATHEDRAL using RMSD, we are not justified in saying RUPEE is better than CATHEDRAL because of the RMSD comparison. However, we can say that RUPEE is better than CATHEDRAL with respect to RMSD and in so far as RMSD is a good measure of structure similarity, this comparison is useful. Below, we make comparisons using TM-score and RMSD in all cases. However, where possible, we also compare to other searches using scoring schemes native to those searches.

In the comparisons for the TM-score, we normalize by either the length of the query structure or the average length of structures being compared. For the TM-score plots, in the vertical axis, we use (q) to indicate normalization by the length of query structure or (avg) to indicate normalization by the average length of the two structures.

### Scoring vs. mTM-align

In Figure 5, we compare the average TM-scores [25] and RMSD scores of the top 100 results for RUPEE and mTM-align [8] using whole PDB chains deposited in the PDB as of 2020-01-01. TM-scores and RMSD scores have been calculated using TM-align [26]. For comparing TM-scores, we use the RUPEE Contained-In search type to search by TM-score normalized by the query structure identical to the scoring used by mTM-align. For comparing RMSD scores, we use the RUPEE RMSD search type. mTM-align does not provide an RMSD search type so when comparing by RMSD we sort the top 100 mTM-align results by RMSD.

**Figure 5.**
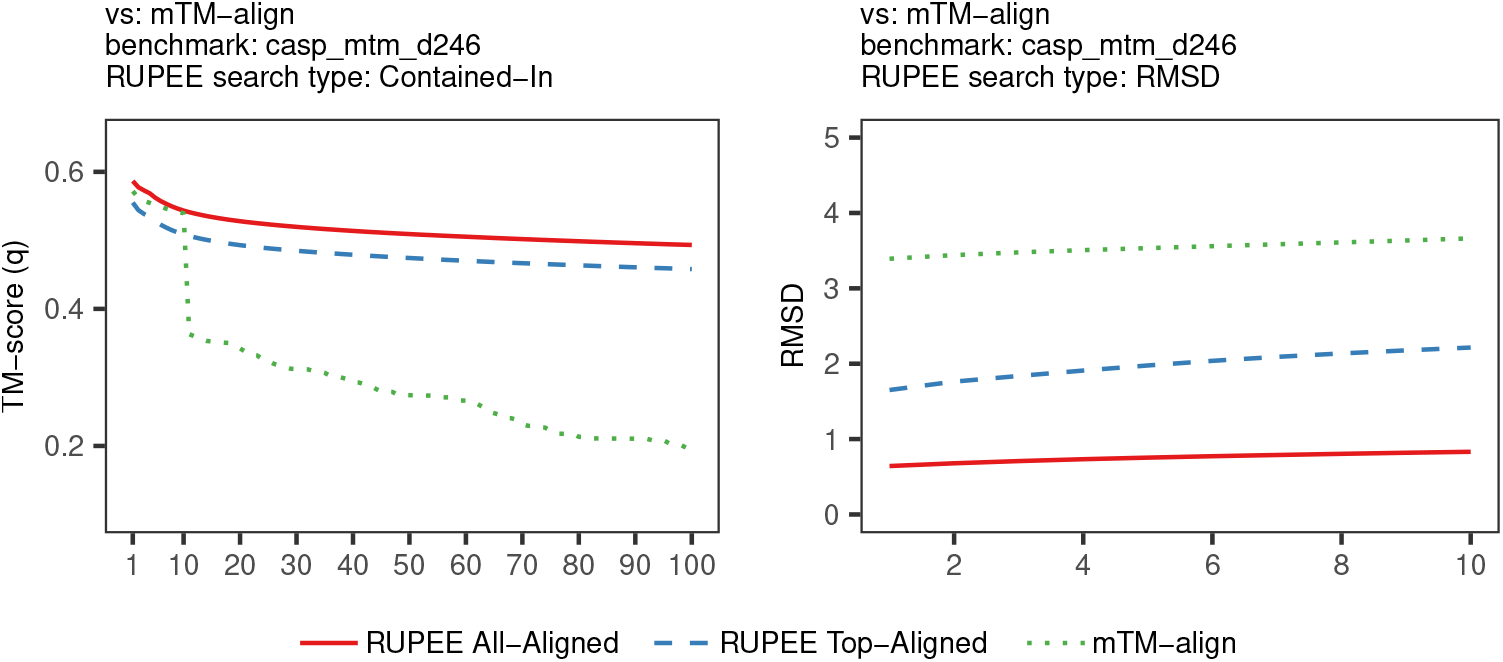
Comparison of average TM-scores and RMSD scores for RUPEE vs. mTM-align

mTM-align only returns 10 results for 95 benchmark structures and returns 100 or more results for only 84 benchmark structures. For TM-score, mTM-align compares favorably with RUPEE all-aligned and top-aligned for the first 10 results, staying within 0.01 TM-score points, but then drops off precipitously. In the left plot of Figure 5 we do not cut off the plot after 10 results because that would be unfair to RUPEE to not highlight the problem that mTM-align has with respect to the number of results it is able to return for difficult searches. For RMSD, both RUPEE all-aligned and top-aligned perform better than mTM-align by a wide margin at all ranks. In the right plot of Figure 5 we do cut off the plot after 10 results because there is no maximum RMSD to let the curve converge to when filling in for missing results. Since protein structure search results usually contain blocks of highly similar structures within the same fold, it is important to return a sufficient number of results in order to show structures from multiple folds in the results, especially when attempting to validate that a predicted structure is within the neighborhood of the expected fold.

### Scoring vs. SSM

In Figure 6, we compare the average Q-scores [9], TM-scores [25] and RMSD scores of the top 100 results for RUPEE and SSM [9] using SCOP v1.73 domains. We calculated TM-scores and Q-scores using TM-align [26] and RMSD scores using the Combinatorial Extension (CE) [29] algorithm. For the TM-score comparison, we normalize by the average length of the compared structures. In addition to TM-score and RMSD, we also compare on Q-score because SSM provides an option to search by Q-score and RMSD. We calculated our own Q-scores and RMSD scores for SSM because we observed the scores they provided are wildly incorrect in many cases. For instance, a large set of results all start with a block of perfect matches with a Q-score of 1.0 and RMSD of 0.0, which is clearly impossible given that we are searching with predicted structures. We did not observe this problem with SSM when searching on known protein structures.

**Figure 6.**
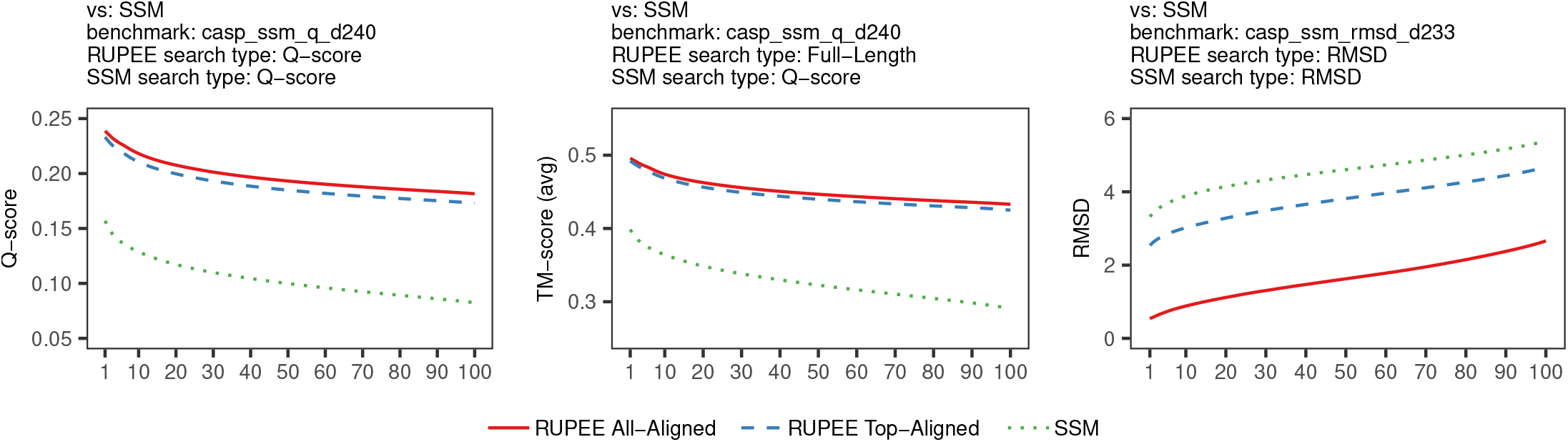
Comparison of average Q-scores, TM-scores and RMSD scores for RUPEE vs. SSM

As shown in Figure 6, both RUPEE all-aligned and top-aligned perform better than SSM at all ranks for Q-scores, TM-scores and RMSD scores. The similarity between the plots for Q-scores and TM-scores suggest some correspondence between how Q-scores and TM-scores are calculated. Both the Q-score and the TM-score are intended as good measures of full-length similarity.

### Scoring vs. CATHEDRAL

In Figure 7, we compare the average SSAP-scores [28], TM-scores [25] and RMSD scores of the top 100 results for RUPEE and CATHEDRAL [10] using CATH v4.2 domains. For comparing to CATHEDRAL, we filter by CATH s35 cluster representatives since that is all that CATHEDRAL returns. For the TM-score comparison, we use TM-align to calculate the scores, and we normalize by the average length of the compared structures. We use the cath-ssap tool provided in the cath-tools suite [30] to calculate SSAP-scores for comparisons. As we do for SSM above, we calculate RMSD scores using CE.

**Figure 7.**
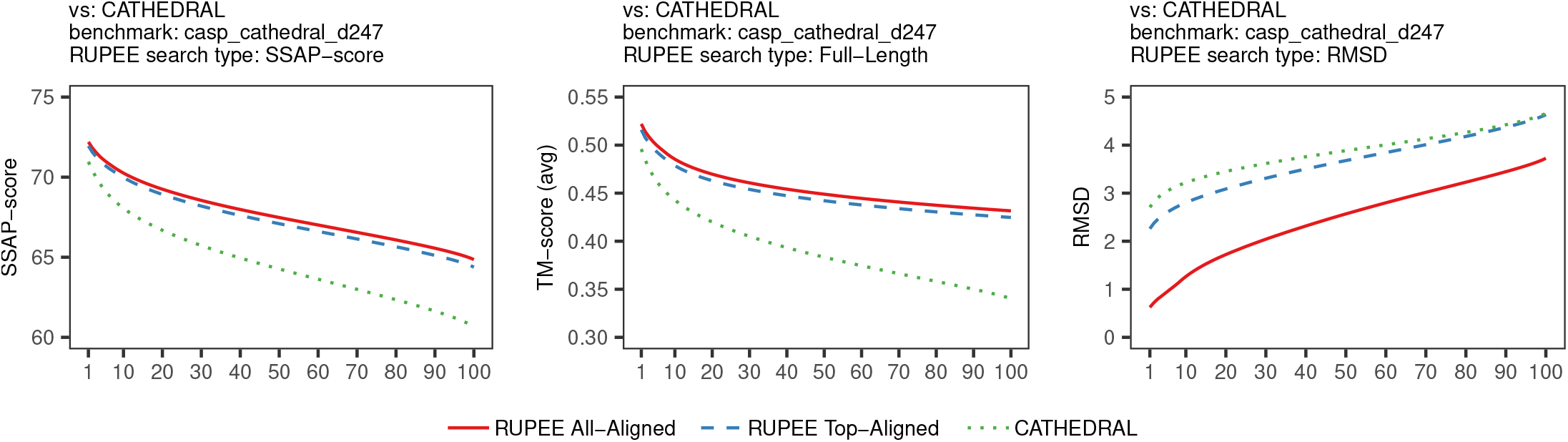
Comparison of average SSAP-scores, TM-scores and RMSD scores for RUPEE vs. CATHEDRAL

As shown in Figure 7, both RUPEE all-aligned and top-aligned perform better than CATHEDRAL at all ranks for SSAP-scores, TM-scores and RMSD scores. It is remarkable that RUPEE performs better than CATHEDRAL using the score that the CATHEDRAL search is based on.

### Scoring vs. VAST

In Figure 8, we compare the average TM-scores and RMSD scores of the top 100 results for RUPEE and VAST. For the TM-score comparison, we use TM-align to calculate the scores, and we normalize by the average length of the compared structures. We were not able to duplicate the VAST-score ourselves for our internal alignment scoring and so do not provide an additional search type as we did for the Q-score and the SSAP-score. As we do for SSM and CATHEDRAL above, we calculate RMSD scores using CE.

**Figure 8.**
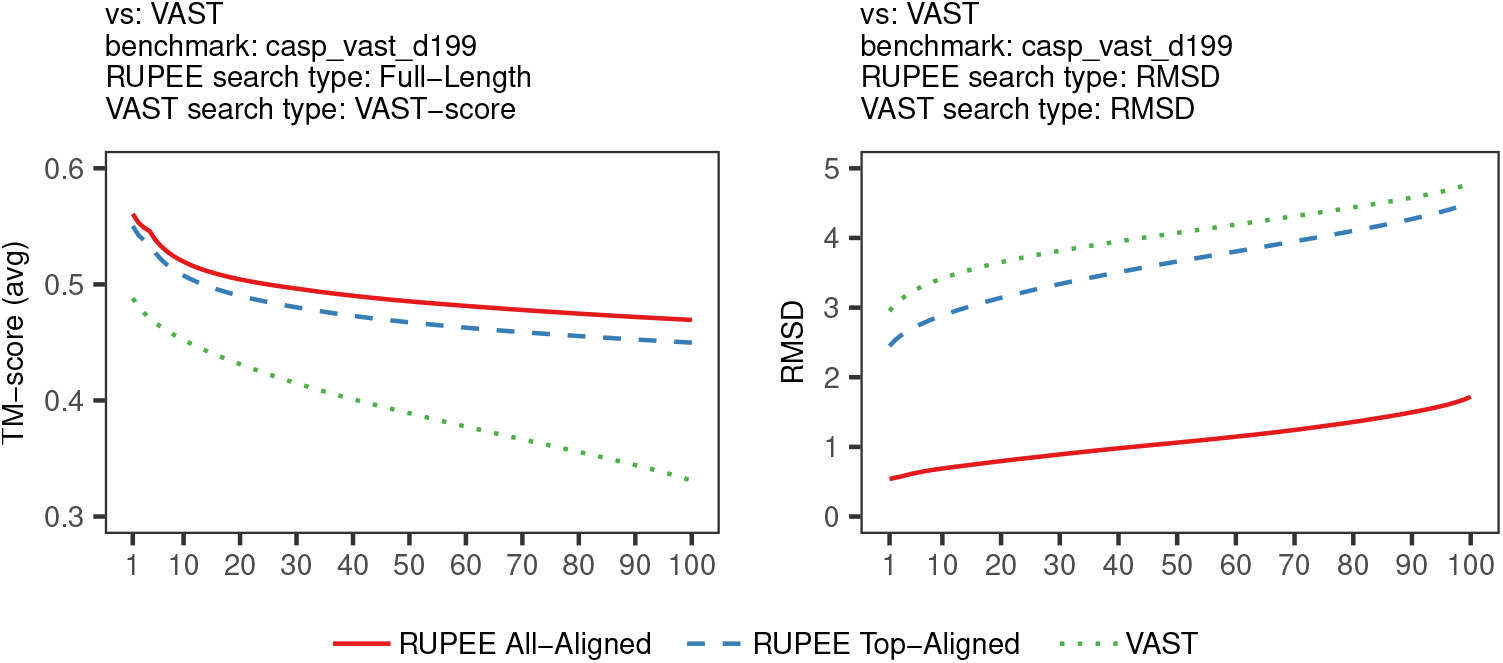
Comparison of average TM-scores and RMSD scores for RUPEE vs. VAST

We run RUPEE against whole PDB chains deposited in the PDB as of 2020-01-01 whereas VAST is run against the molecular modelling database (MMDB), which augments familiar PDB structures with knowledge annotations, although it is not clear what version of the PDB they are operating with. Despite possible differences in the structural databases searched, it is the case that all the VAST structure results for our benchmark do match to structures contained in the PDB as of 2020-01-01.

As shown in Figure 8, both RUPEE all-aligned and top-aligned perform better than VAST at all ranks for TM-scores and RMSD scores. In the left plot of Figure 8, for the TM-score comparison, we use the VAST-score sort provided by VAST and in the right plot, for the RMSD comparison, we use the RMSD score sort provided by VAST when collecting the data and then sort the top 100 results based on the compared score as usual. Since the VAST-score is roughly a full length score, we use the TM-score normalized by the average length of both structures for comparisons.

### Scoring all-aligned vs. top-aligned

For full length searches, in Figure 9, we compare the results of RUPEE all-aligned to RUPEE top-aligned for two different benchmarks against the same structure database, SCOP v2.07, consisting of more than 250,000 structures. The easier benchmark, scop_d360, is the benchmark of known proteins structures that we used in our previous work on RUPEE [7]. The harder benchmark, casp_d250, is the benchmark of protein structure predictions from above. For scop_d360, the difference between RUPEE all-aligned and top-aligned is only a fraction of a TM-score point, whereas for casp_d250, all-aligned is 0.01 to 0.02 TM-score points better than top-aligned across all ranks except for the first 10. Figure 9 suggest that RUPEE all-aligned is more suitable than RUPEE top-aligned for searching on protein structure predictions. However, for known protein structures, the performance of top-aligned is almost identical to all-aligned.

**Figure 9.**
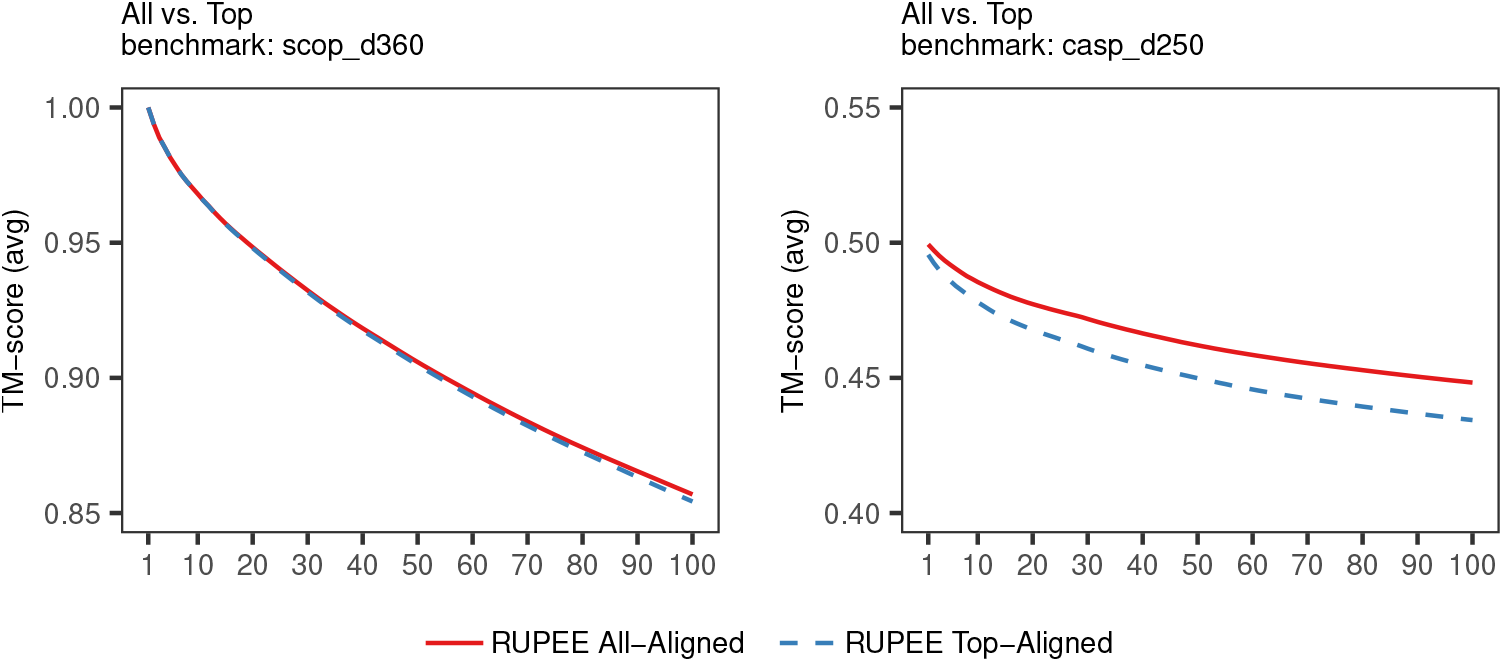
RUPEE vs RUPEE. Comparison of average TM-scores provided by TM-align for top 100 results ranked by TM-score

For RMSD searches, the right plots of Figs 5, 6, 7 and 8 all shows RUPEE all-aligned doing significantly better than top-aligned across all ranks. This is notably different from Figure 9. Part of the reason for this larger difference is that all-aligned mode starts filtering results on the compared score much earlier in the pipeline whereas top-aligned mode starts filtering results on the compared score only after the initial filtering and NW alignments.

### Sample Alignments

Figure 10 shows the structure alignments of the top-scoring full-length structures matches that RUPEE identified that none of the other protein structure searches we compared to identified. While we recognize that mTM-align, SSM, CATHEDRAL and VAST may have performed equal to or better than RUPEE on some benchmark structures, RUPEE performs better on average as was shown in Figs 5, 6, 7 and 8.

**Figure 10.**
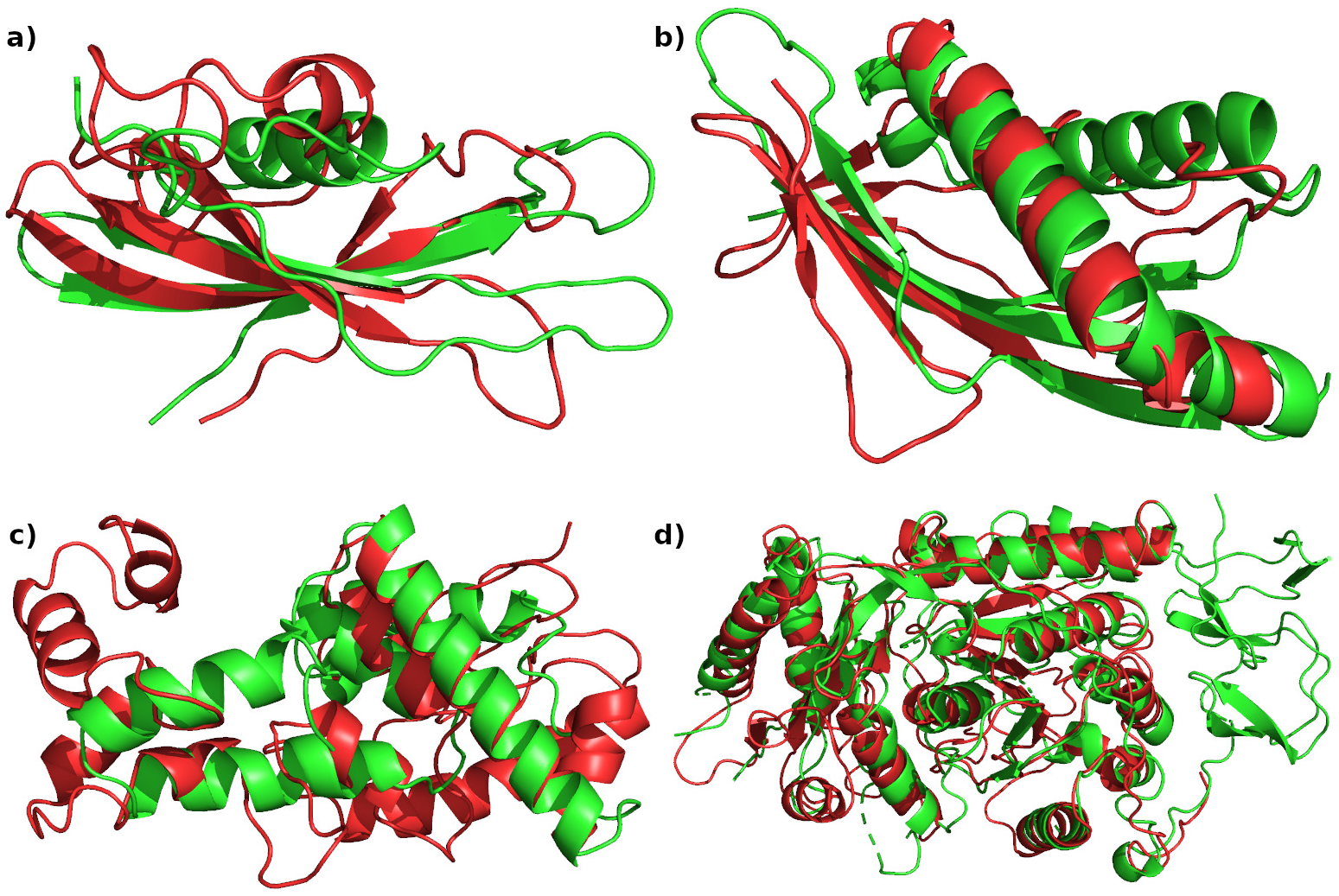
Red structures are CASP predictions. (a) T0960TS354-D2 aligned with 2lyxA00 found in CATH v4.2. (b) T0980s1TS043-D1 aligned with 6ahqC found among whole PDB chains. (c) T0990TS089-D3 aligned with 1w07A01 found in CATH v4.2. (d) T1000TS043-D2 aligned with 6u7lD found among whole PDB chains.

Figure 10 illustrates qualitatively the types of difficult matches RUPEE has no problem with finding despite the complexity of the loops. We believe it is these types of difficult matches that would be of interest to researchers investigating protein structure prediction.

## Conclusion

With the recent successes of coevolutionary methods at CASP11 [5] and deep learning with AlpahFold at CASP13 [3], the long-standing problem of protein structure prediction has seen renewed interest. Despite this renewed interest, the problem of identifying the most similar known protein structures to structure predictions has not been explored and researchers are left navigating a variety of structure searches not specifically designed for this purpose. In addition to matching the performance of some of the best available protein structure searches on a benchmark of known protein structures as shown in our previous work on RUPEE [7], we have now shown that RUPEE effectively addresses the problem of searching on structure predictions and is uniquely suited to support the development of protein structure prediction methods.

## Supporting information

S1 Benchmarks

## Supporting Information

**S1 Benchmarks. Structures included in benchmarks used for evaluation.**

## References

1. Burley SK, Berman HM, Christie C, Duarte JM, Feng Z, Westbrook J, et al. RCSB Protein Data Bank: Sustaining a living digital data resource that enables breakthroughs in scientific research and biomedical education. Protein Science. 2018;27(1):316–330.

2. Consortium TU. UniProt: The universal protein knowledgebase. Nucleic Acids Research. 2017;45(D1):D158–D169.

3. AlQuraishi M. AlphaFold at CASP13. Bioinformatics. 2019;.

4. Moult J, Pedersen JT, Judson R, Fidelis K. A large-scale experiment to assess protein structure prediction methods. Proteins: Structure, Function, and Bioinformatics. 1995;23(3):ii–iv.

5. Moult J, Fidelis K, Kryshtafovych A, Schwede T, Tramontano A. Critical assessment of methods of protein structure prediction: Progress and new directions in round XI. Proteins: Structure, Function, and Bioinformatics. 2016;84(S1):4–14.

6. Rohl CA, Strauss CEM, Misura KMS, Baker D. Protein Structure Prediction Using Rosetta. In: Numerical Computer Methods, Part D. vol. 383 of Methods in Enzymology. Academic Press; 2004. p. 66–93.

7. Ayoub R, Lee Y. RUPEE: A fast and accurate purely geometric protein structure search. PLOS ONE. 2019;14(3):1–17.

8. Dong R, Pan S, Peng Z, Zhang Y, Yang J. mTM-align: a server for fast protein structure database search and multiple protein structure alignment. Nucleic Acids Research. 2018;46(July):380–386.

9. Krissinel E, Henrick K. Secondary-structure matching (SSM), a new tool for fast protein structure alignment in three dimensions. Acta Crystallographica Section D: Biological Crystallography. 2004;60(12 I):2256–2268.

10. Redfern OC, Harrison A, Dallman T, Pearl FMG, Orengo CA. CATHEDRAL: A Fast and Effective Algorithm to Predict Folds and Domain Boundaries from Multidomain Protein Structures. PLoS Comput Biol. 2007;3(11):e232.

11. Chothia C, Lesk AM. The relation between the divergence of sequence and structure in proteins. The EMBO Journal. 1986;5(4):823–826.

12. Illergård K, Ardell DH, Elofsson A. Structure is three to ten times more conserved than sequence—A study of structural response in protein cores. Proteins: Structure, Function, and Bioinformatics. 2009;77(3):499–508.

13. Gibrat JF, Madej T, Bryant SH. Surprising similarities in structure comparison. Current Opinion in Structural Biology. 1996;6(3):377–385.

14. Kinch LN, Kryshtafovych A, Monastyrskyy B, Grishin NV. CASP13 target classification into tertiary structure prediction categories. Proteins: Structure, Function, and Bioinformatics. 2019;87(12):1021–1036.

15. Orengo C, Mitchie A, Jones S, Jones DT, Swindells M, Thornton JM. CATH - A hierarchic classification of protein domain structures. Structure. 1997;5(8):1093–1109.

16. Fox NK, Brenner SE, Chandonia JM. SCOPe: Structural Classification of Proteins - extended, integrating SCOP and ASTRAL data and classification of new structures. Nucleic Acids Res. 2014;42(1):304–309.

17. Cheng H, Schaeffer RD, Liao Y, Kinch LN, Pei J, Shi S, et al. ECOD: An Evolutionary Classification of Protein Domains. PLoS Computational Biology. 2014;10(12).

18. Kabsch W, Sander C. Dictionary of protein secondary structure: Pattern recognition of hydrogen-bonded and geometrical features. Biopolymers. 1983;22(12):2577–2637.

19. Broder AZ. On the resemblance and containment of documents. In: Proc. Compression and Complexity of Sequences. Positano, Italy; 1997. p. 21–29.

20. Levandowsky M, Winter D. Distance between Sets. Nature. 1971;234(5323):34–35.

21. Broder AZ, Charikar M, Frieze AM, Mitzenmacher M. Min-wise independent permutations. In: ACM Symposium on Theory of Computing. Dallas, USA; 1998. p. 327–336.

22. Indyk P, Motwani R. Approximate nearest neighbors: towards removing the curse of dimensionality. In: ACM Symposium on Theory of Computing. Dallas, USA; 1998. p. 604–613.

23. Needleman SB, Wunsch CD. A general method applicable to the search for similarities in the amino acid sequence of two proteins. Journal of Molecular Biology. 1970;48(3):443–453.

24. Brudno M, Malde S, Poliakov A, Do CB, Couronne O, Dubchak I, et al. Glocal alignment: finding rearrangements during alignment. Bioinformatics. 2003;19:i54–i62.

25. Zhang Y, Skolnick J. Scoring function for automated assessment of protein structure template quality. Proteins. 2004;57(4):702–710.

26. Zhang Y, Skolnick J. TM-align: A protein structure alignment algorithm based on the TM-score. Nucleic Acids Research. 2005;33(7):2302–2309.

27. Holm L, Rosenström P. Dali server: Conservation mapping in 3D. Nucleic Acids Research. 2010;38(SUPPL. 2):1–5.

28. Orengo CA. Protein Structure Alignment. J Mol Biol. 1989;.

29. Shindyalov IN, Bourne PE. Protein structure alignment by incremental combinatorial extension (CE) of the optimal path. Protein Eng Des Sel. 1998;11(9):739–747.

30. Dawson NL, Lewis TE, Das S, Lees JG, Lee D, Ashford P, et al. CATH: an expanded resource to predict protein function through structure and sequence. Nucleic Acids Res. 2017;45(D1):D289–D295.

