## Supplementary material for "Protein structure search to support the development of protein structure prediction methods": S1 Benchmarks

### 1 Benchmarks

#### 1.1 casp_d250

T0953s1TS043-D1
T0953s1TS089-D1
T0953s1TS145-D1
T0953s1TS196-D1
T0953s1TS197-D1
T0953s1TS224-D1
T0953s1TS261-D1
T0953s1TS322-D1
T0953s1TS354-D1
T0953s1TS498-D1
T0957s2TS043-D1
T0957s2TS089-D1
T0957s2TS145-D1
T0957s2TS196-D1
T0957s2TS197-D1
T0957s2TS224-D1
T0957s2TS261-D1
T0957s2TS322-D1
T0957s2TS354-D1
T0957s2TS498-D1
T0960TS043-D2
T0960TS089-D2
T0960TS145-D2
T0960TS196-D2
T0960TS197-D2
T0960TS224-D2
T0960TS261-D2
T0960TS322-D2
T0960TS354-D2
T0960TS498-D2
T0963TS043-D2
T0963TS089-D2
T0963TS145-D2
T0963TS196-D2
T0963TS197-D2
T0963TS224-D2
T0963TS261-D2
T0963TS322-D2
T0963TS354-D2
T0963TS498-D2
T0968s1TS043-D1
T0968s1TS089-D1

T0968s1TS145-D1
T0968s1TS196-D1
T0968s1TS197-D1
T0968s1TS224-D1
T0968s1TS261-D1
T0968s1TS322-D1
T0968s1TS354-D1
T0968s1TS498-D1
T0968s2TS043-D1
T0968s2TS089-D1
T0968s2TS145-D1
T0968s2TS196-D1
T0968s2TS197-D1
T0968s2TS224-D1
T0968s2TS261-D1
T0968s2TS322-D1
T0968s2TS354-D1
T0968s2TS498-D1
T0969TS043-D1
T0969TS089-D1
T0969TS145-D1
T0969TS196-D1
T0969TS197-D1
T0969TS224-D1
T0969TS261-D1
T0969TS322-D1
T0969TS354-D1
T0969TS498-D1
T0975TS043-D1
T0975TS089-D1
T0975TS145-D1
T0975TS196-D1
T0975TS197-D1
T0975TS224-D1
T0975TS261-D1
T0975TS322-D1
T0975TS354-D1
T0975TS498-D1
T0980s1TS043-D1
T0980s1TS089-D1
T0980s1TS145-D1
T0980s1TS196-D1
T0980s1TS197-D1
T0980s1TS224-D1
T0980s1TS261-D1
T0980s1TS322-D1

T0980s1TS354-D1
T0980s1TS498-D1
T0986s2TS043-D1
T0986s2TS089-D1
T0986s2TS145-D1
T0986s2TS196-D1
T0986s2TS197-D1
T0986s2TS224-D1
T0986s2TS261-D1
T0986s2TS322-D1
T0986s2TS354-D1
T0986s2TS498-D1
T0987TS043-D1
T0987TS043-D2
T0987TS089-D1
T0987TS089-D2
T0987TS145-D1
T0987TS145-D2
T0987TS196-D1
T0987TS196-D2
T0987TS197-D1
T0987TS197-D2
T0987TS224-D1
T0987TS224-D2
T0987TS261-D1
T0987TS261-D2
T0987TS322-D1
T0987TS322-D2
T0987TS354-D1
T0987TS354-D2
T0987TS498-D1
T0987TS498-D2
T0989TS043-D1
T0989TS043-D2
T0989TS089-D1
T0989TS089-D2
T0989TS145-D1
T0989TS145-D2
T0989TS196-D1
T0989TS196-D2
T0989TS197-D1
T0989TS197-D2
T0989TS224-D1
T0989TS224-D2
T0989TS261-D1
T0989TS261-D2

T0989TS322-D1
T0989TS322-D2
T0989TS354-D1
T0989TS354-D2
T0989TS498-D1
T0989TS498-D2
T0990TS043-D1
T0990TS043-D3
T0990TS089-D1
T0990TS089-D3
T0990TS145-D1
T0990TS145-D3
T0990TS196-D1
T0990TS196-D3
T0990TS197-D1
T0990TS197-D3
T0990TS224-D1
T0990TS224-D3
T0990TS261-D1
T0990TS261-D3
T0990TS322-D1
T0990TS322-D3
T0990TS354-D1
T0990TS354-D3
T0990TS498-D1
T0990TS498-D3
T0998TS043-D1
T0998TS089-D1
T0998TS145-D1
T0998TS196-D1
T0998TS197-D1
T0998TS224-D1
T0998TS261-D1
T0998TS322-D1
T0998TS354-D1
T0998TS498-D1
T1000TS043-D2
T1000TS089-D2
T1000TS145-D2
T1000TS196-D2
T1000TS197-D2
T1000TS224-D2
T1000TS261-D2
T1000TS322-D2
T1000TS354-D2
T1000TS498-D2

T1001TS043-D1
T1001TS089-D1
T1001TS145-D1
T1001TS196-D1
T1001TS197-D1
T1001TS224-D1
T1001TS261-D1
T1001TS322-D1
T1001TS354-D1
T1001TS498-D1
T1010TS043-D1
T1010TS089-D1
T1010TS145-D1
T1010TS196-D1
T1010TS197-D1
T1010TS224-D1
T1010TS261-D1
T1010TS322-D1
T1010TS354-D1
T1010TS498-D1
T1015s1TS043-D1
T1015s1TS089-D1
T1015s1TS145-D1
T1015s1TS196-D1
T1015s1TS197-D1
T1015s1TS224-D1
T1015s1TS261-D1
T1015s1TS322-D1
T1015s1TS354-D1
T1015s1TS498-D1
T1017s2TS043-D1
T1017s2TS089-D1
T1017s2TS145-D1
T1017s2TS196-D1
T1017s2TS197-D1
T1017s2TS224-D1
T1017s2TS261-D1
T1017s2TS322-D1
T1017s2TS354-D1
T1017s2TS498-D1
T1021s3TS043-D1
T1021s3TS043-D2
T1021s3TS089-D1
T1021s3TS089-D2
T1021s3TS145-D1
T1021s3TS145-D2

T1021s3TS196-D1
T1021s3TS196-D2
T1021s3TS197-D1
T1021s3TS197-D2
T1021s3TS224-D1
T1021s3TS224-D2
T1021s3TS261-D1
T1021s3TS261-D2
T1021s3TS322-D1
T1021s3TS322-D2
T1021s3TS354-D1
T1021s3TS354-D2
T1021s3TS498-D1
T1021s3TS498-D2
T1022s1TS043-D1
T1022s1TS089-D1
T1022s1TS145-D1
T1022s1TS196-D1
T1022s1TS197-D1
T1022s1TS224-D1
T1022s1TS261-D1
T1022s1TS322-D1
T1022s1TS354-D1
T1022s1TS498-D1

#### 1.2 casp_mtm_d246

Same as casp_d250 excluding:

T0957s2TS354-D1
T0960TS089-D2
T0975TS145-D1
T0975TS196-D1

#### 1.3 casp_ssm_q_d240

Same as casp_d250 excluding:

T0953s1TS043-D1
T0953s1TS089-D1
T0957s2TS145-D1
T0968s2TS197-D1
T0989TS043-D2
T1010TS089-D1
T1010TS322-D1
T1015s1TS197-D1

T1015s1TS261-D1
T1017s2TS145-D1

##### 1.4 casp_ssm_rmsd_d233

Same as casp_d250 excluding:

T0953s1TS043-D1
T0953s1TS089-D1
T0957s2TS197-D1
T0968s1TS322-D1
T0968s2TS197-D1
T0989TS261-D1
T1000TS145-D2
T1000TS224-D2
T1000TS261-D2
T1000TS322-D2
T1000TS354-D2
T1001TS197-D1
T1001TS224-D1
T1010TS089-D1
T1010TS322-D1
T1015s1TS043-D1
T1017s2TS089-D1

##### 1.5 casp_cathedral_d247

Same as casp_d250 excluding:

T0953s1TS498-D1
T0969TS498-D1
T0980s1TS498-D1

##### 1.6 casp_vast_d199

Same as casp_d250 excluding:

T0953s1TS498-D1
T0968s1TS196-D1
T0969TS196-D1
T0969TS354-D1
T0975TS043-D1
T0975TS089-D1
T0975TS196-D1
T0975TS261-D1
T0975TS322-D1
T0975TS354-D1

T0975TS498-D1
T0980s1TS224-D1
T0980s1TS261-D1
T0980s1TS322-D1
T0986s2TS354-D1
T0987TS089-D2
T0987TS145-D2
T0987TS261-D2
T0987TS322-D2
T0987TS354-D2
T0987TS498-D2
T0989TS145-D2
T0989TS196-D2
T0990TS043-D3
T0990TS089-D1
T0990TS089-D3
T0990TS145-D1
T0990TS196-D1
T0990TS196-D3
T0990TS197-D1
T0990TS224-D1
T0990TS224-D3
T0990TS261-D1
T0990TS261-D3
T0990TS322-D1
T0990TS322-D3
T0990TS354-D1
T0990TS498-D3
T0998TS043-D1
T0998TS089-D1
T0998TS145-D1
T0998TS196-D1
T0998TS261-D1
T0998TS322-D1
T0998TS354-D1
T0998TS498-D1
T1010TS043-D1
T1010TS089-D1
T1010TS261-D1
T1010TS354-D1
T1021s3TS043-D1
